## Supplementary Information for "Within-host competition modulates pneumococcal antibiotic resistance in the pre-vaccination era"

### Model description and data fitting

#### Epidemiological framework

We model the transmission dynamics of pneumococcal strains according to the epidemiological framework by Obolski et al. (Obolski et al. 2018). We define pneumococcal strains by the combination of their serotype and antibiotic resistance profile to two distinct antibiotics. Similarly to the nomenclature used in previous studies, each strain's genotype is conceptualised by a tuple $\{i,j\}$, where $i$ determines the serotype and $j$ the antibiotic resistance profile. We model two serotypes only, such that $i \in \{a,b\}$. For the entirety of our study, we define serotype $a$ as a vaccine type (VT) and serotype $b$ as a non-vaccine type (NVT). We also model two antibiotics $x$ and $y$ and consider only resistance to each independently, with $j \in \{00, 01, 10\}$, where $j = 00$ refers to sensitive strains, $j = 01$ to strains resistant only to antibiotic $y$, and $j = 10$ to strains resistant only to antibiotic $x$. For instance, a strain defined by $i = a, j = 00$ is of VT serotype $a$ and is sensitive to both antibiotics; while a strain $i = b, j = 01$ is of NVT serotype $b$, sensitive to antibiotic $x$ and resistant to antibiotic $y$.

We keep the terminology of the original model formulation regarding host epidemiological states, thus denoting $y_{i,j}$ as the proportion of individuals carrying strain $i,j$, and $Z_i$ the proportion of the population previously exposed to serotype $i$. Other relevant states include: $S_i$ as the proportion of the population naive to serotype $i$; $y_i$ as the proportion carrying strains of serotype $i$ (independently of resistance $j$); and $y_j$ as the proportion carrying strains with a resistance profile $j$ (independently of serotype $i$). We also keep the terminology for the epidemiological parameters: $\gamma$ is serotype-specific immunity; $\psi$ is the probability that a carried susceptible strain ($j = 00$) will suppress host co-colonization by a resistant strain ($j = 01$ or $j = 10$) due to the fitness cost of antibiotic resistance (details below); $1/\mu$ is the host life-span; $1/\sigma$ is the host carriage duration; R0 is the basic reproduction number and $\beta$ is the transmission rate.

Individuals are born naive to all strains, with the size of the host population kept constant (deaths being replaced by newborns). Colonization results in carriage for an average duration of $1/\sigma$ days, from which recovery (clearance) may lead to complete serotype immunity if $\gamma = 1$ or partial life-long immunity if $\gamma < 1$. Thus, if $\gamma < 1$, hosts can be re-colonized by the same serotype throughout their lifetime, with increasing likelihood as $\gamma$ tends to zero. Co-colonization of up to two strains is allowed, unless $\gamma = 1$ and strains belong to the same serotype. Within-host strain competition interferes with co-colonization if $\psi > 0$, which captures the degree to which intrinsic fitness differences (e.g. growth rates) between resistant and sensitive strains may allow a currently carried sensitive strain ($j = 00$) to suppress co-colonization by a resistant strain ($j = 01$, $j = 10$). We emphasize that $\psi$ models a form of competition between bacterial strains that is not mediated by immunity - from now onwards referred to as ecological competition. The modelled processes related to an individual's colonization and epidemiological states are summarised in **Figure 1A** in the main text.

The intrinsic transmissibility of each strain is defined by a basic reproductive number $R0_{i,j} = \beta_{i,j} / (\sigma_{i,j} + \mu_{i,j})$ . As in the original framework, we assume that the fitness cost of antibiotic resistance can translate into lower infectivity of resistant strains ($\beta_{i,j=00} > \beta_{i,j=01}$ or $\beta_{i,j=00} > \beta_{i,j=10}$ ). However, due to antibiotic usage, the duration of carriage may be longer for resistant strains ( $1/\sigma_{i,j=10} > 1/\sigma_{i,j=00}$ or $1/\sigma_{i,j=01} > 1/\sigma_{i,j=00}$ ) (Lehtinen et al. 2017). Thus, in the absence of antibiotic usage, the $R_0$ of resistant strains ( $R0_{i,j=01}$ , $R0_{i,j=10}$ ) will typically be lower than the $R0$ of sensitive strains ( $R0_{i,j=00}$ ), but this is likely reversed with antibiotic usage. We define $\Delta_{i,j}$ as the ratio of $R0$ of a resistant strain ( $i,j = 01$ or $i,j = 10$ ) compared to a sensitive strain ( $i,j = 00$ ) (e.g. $R0^{\,i,j=01} = \Delta_{i,j} \times R0^{\,i,j=00}$ ).

#### Model parameterization

Unless stated otherwise, we assume the default parameters to be: $\eta = 0.033$ (average introduction of one infection per month), $\omega = 0.0001$ , $R_0^{VT} = 2.5$ and $R_0^{NVT} = 2.0$ (Nurhonen, Cheng, and Auranen 2013; le Polain de Waroux et al. 2018; Hoti et al. 2009), $\sigma = 0.033$ (average infectious period of 30 days (Högberg et al. 2007), see Table S1 of (Lourenço et al. 2019) for literature review on this parameter), $1/\mu = 75$ (average life-span of 75 years), $\Delta_{i,j=01} = 1$ and $\Delta_{i,j=10} = 1$ (i.e. no difference in the $R_0$ of sensitive and resistant strains), $N \approx 1$ million (host-population size), and $LxL = 529$ (host-population structure 23 x 23 with communities of size ~1890 individuals). In sensitivity exercises, we model parameter variations such as population structure $LxL$ , $R_0^{VT}$ and $R_0^{NVT}$ and host mobility $\omega$ (which are presented in **Supplementary Figures** as referenced throughout the main text).

#### Data sources and findings

Data source used was the European Centre for Disease Prevention and Control, from the surveillance atlas (antimicrobial resistance) for the year 2005, vastly representative of a pre-PCV era across European countries (available at ("European Centre for Disease Control Antibiotic Resistance Data" n.d.)). The year 2005 is the earliest made available by the ECDC. The data variables used were (i) the percent of samples sensitive to penicillins and macrolides (independently), and (ii) consumption defined by daily doses per 1000 people per day. We considered solely countries which had a number of samples larger than 50, which included 19 countries. Resistance level was aggregated across countries (EU) using a logistic regression model with random-effect intercepts for each country. Raw data on resistance per country and aggregated EU is presented in **Supplementary Figures S1-2**; and for consumption in **Supplementary Figure S3**.

#### Approximate Bayesian computation

For target (i), we used ECDC data as described in the above sections. For target (ii) we created a dataset of frequencies of VT and NVT samples from 6 studies reporting from before PCV-7 introduction (Syrjänen et al. 2001; Regev-Yochay et al. 2004; Bogaert et al. 2001; Sener et al. 1998; Hussain et al. 2005; Meats et al. 2003). From these pooled frequencies, we sampled, with replacement, 100,000 replicates of frequencies of VTs and of NVTs. These were used to get a distribution of the observed, cross-country relative NVT frequency in the population (**Supplementary Figure S4**). The median of this frequency (~0.15) was compared to the analogous output from the simulations. Finally, for target (iii) we compared model co-infection results with a value of 0.2 and 0.3 (equating to 20 or 30% of carriers being co-infected). We note that observed levels of pneumococcal co-infection are generally higher than this threshold (Tabatabaei et al. 2014; Kamng'ona et al. 2015), but a direct comparison to our model is not possible - we model 2 classes of serotypes ($i \in \{a,b\}$) with 3 genotypes each ($j \in \{00,01,10\}$), which cannot result in levels of co-infection similar to those observed in the real host-pathogen system which is composed of >100 serotypes and many more genotypes. Given a lack of data support for the pneumococcus, we chose a value of 0.2 to represent feasible co-infection levels between resistant and susceptible strains as observed for *Staphylococcus aureus* (Mongkolrattanothai et al. 2011).

### Supplementary Figures

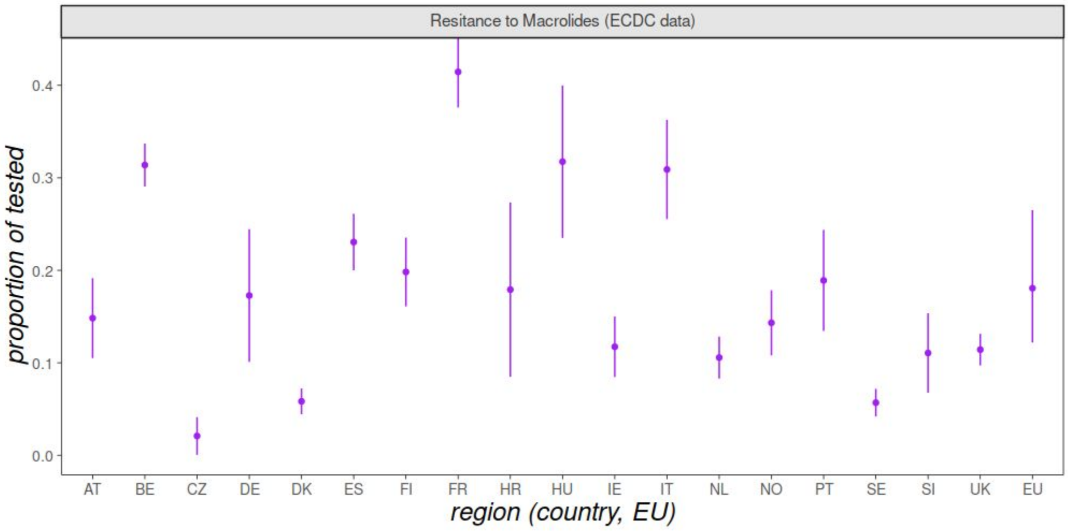


**Figure S1 - ECDC resistance data for macrolides per country and aggregated estimation for EU.** Resistance level to Macrolides (proportion of samples sensitive to Macrolides) for the year 2005 across European countries: AT = Austria, BE = Belgium, CZ = Czechia, DE = Germany, DK = Denmark, ES = Spain, FI = Finland, FR = France, HR = Croatia, HU = Hungary, IE = Ireland, IT = Italy, NL = Netherlands, NO = Norway, PT = Portugal, SE = Sweden, SI = Slovenia, UK = United Kingdom. Countries presented are restricted to those with N>50 samples in 2005. EU is the estimated, aggregated resistance level for the European region as described in the main text.

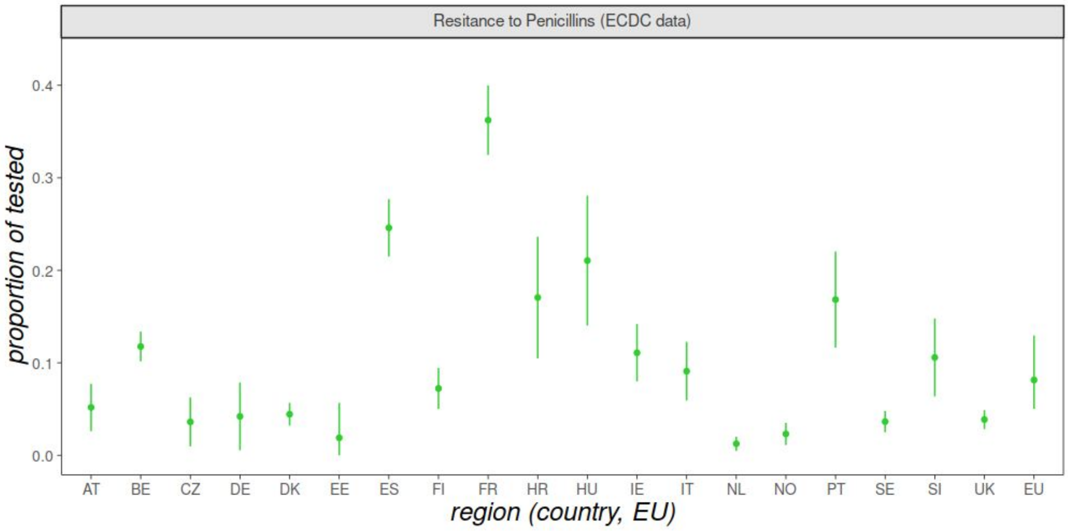

**Figure S2 - ECDC resistance data for penicillins per country and aggregated estimation for EU.** Legend the same as Figure S1.

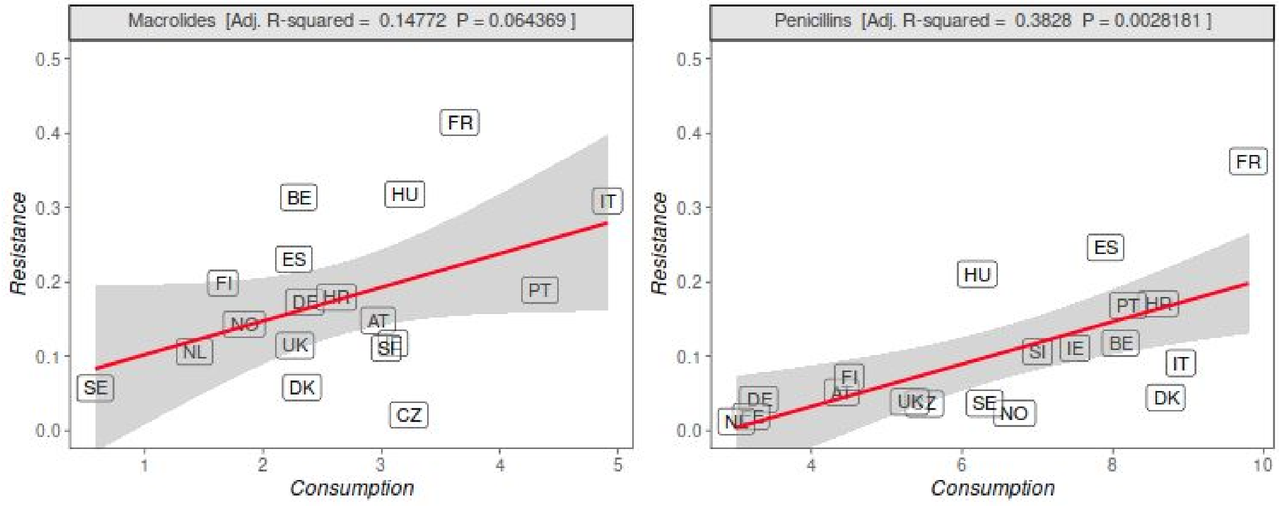


**Figure S3 - ECDC resistance versus consumption data per country. (A)** Resistance level to Macrolides (proportion of samples sensitive to Macrolides) for the year 2005 across European countries: AT = Austria, BE = Belgium, CZ = Czechia, DE = Germany, DK = Denmark, ES = Spain, FI = Finland, FR = France, HR = Croatia, HU = Hungary, IE = Ireland, IT = Italy, NL = Netherlands, NO = Norway, PT = Portugal, SE = Sweden, SI = Slovenia, UK = United Kingdom. Countries presented are restricted to those with N>50 samples in 2005. **(B)** Same as panel A for Penicillins. **(A-B)** Consumption is defined by the ECDC as defined daily doses (DDDs) per 1000 people per day.

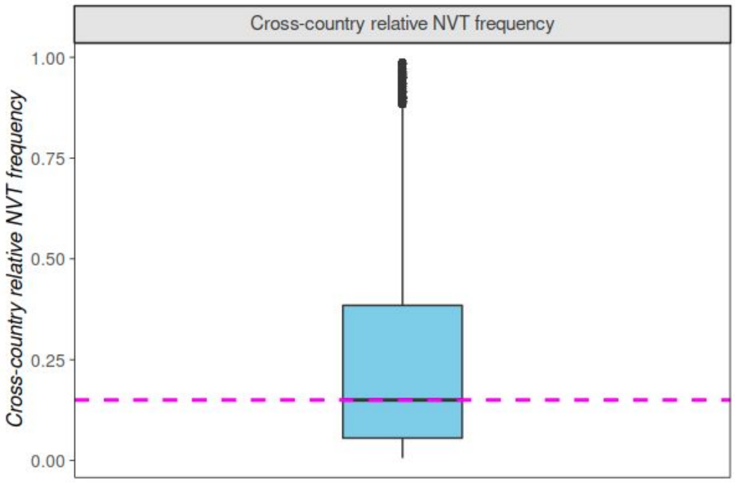


**Figure S4 - Cross-country relative NVT frequency in the population.** Frequencies of VT and NVT samples from 6 studies reporting carriage from before PCV-7 introduction were used to obtain frequencies of VT / (VT + NVT), by sampling with replacement 100,000 replicates of frequencies of VTs and of NVTs. The boxplot is the resulting distribution. The magenta dashed line is the median obtained of ~0.15. Countries sampled in the studies included: Finland, Israel, UK, the Netherlands, and Turkey. See Methods in the main text for details.

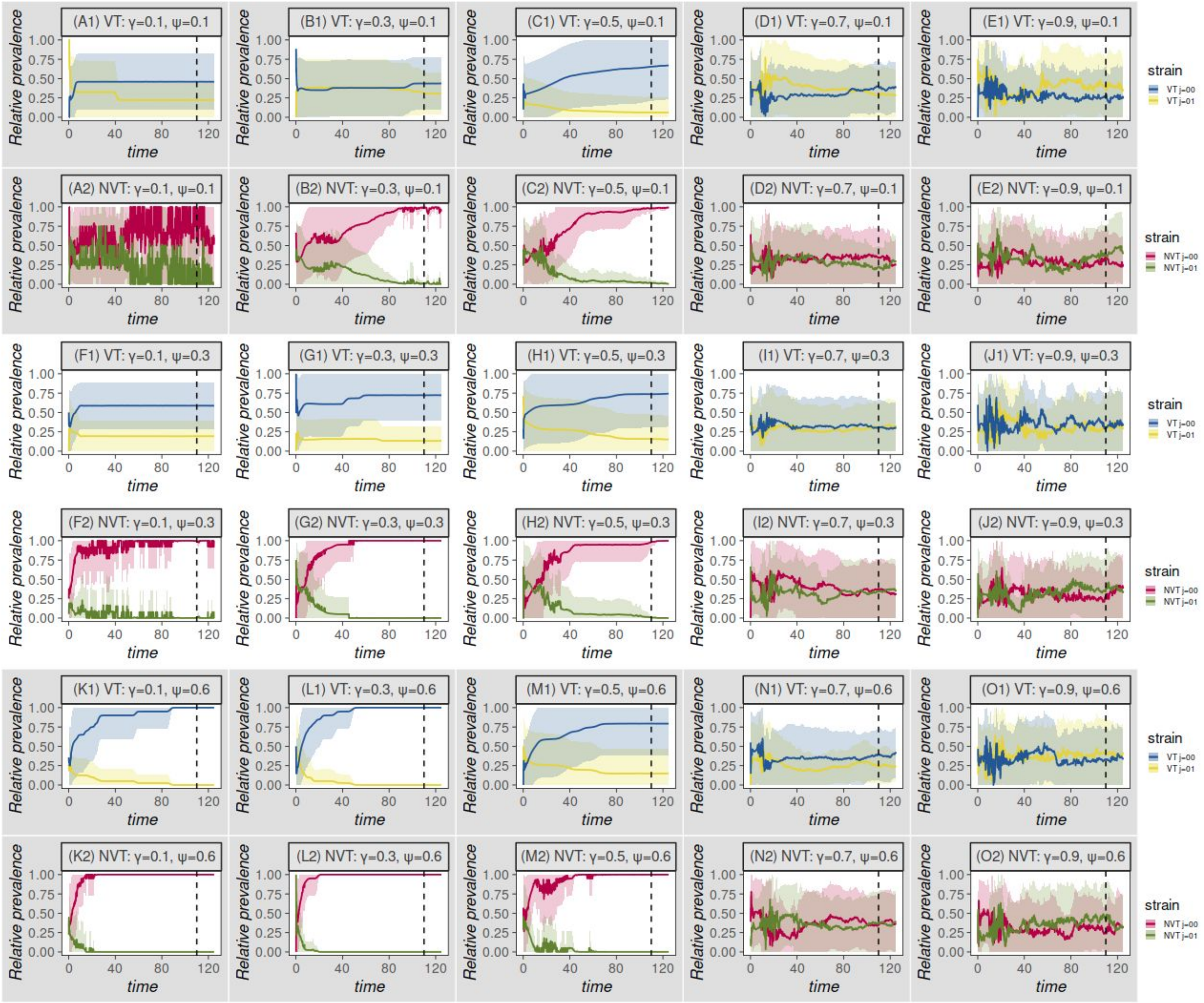


**Figure S5 - Examples of model output with differences in R0 between VT and NVT.** Examples of dynamic output (simulations N=20) when varying the competition parameters (ecological $\psi$, immunological $\gamma$, values show in each panel's title). Output is shown for susceptible ($j = 00$) and one resistant strain ($j = 01$) for both vaccine (VT) and non-vaccine (NVT) types. First row (grey background) includes output for $\psi = 0.2$ and varying $\gamma$. Second row (white background) includes output for $\psi = 0.4$ and varying $\gamma$. Third row (grey background) includes output for $\psi = 0.7$ and varying $\gamma$. For each row, VT dynamics are presented on the top and NVT on the bottom. Lines are the mean dynamic output and shaded areas the standard deviation. Dashed vertical line marks what could be the assumed start of equilibrium. Different parameter combinations would require different waiting times to reach equilibrium. Modelled differences in $R_0$ were: $R_0^{VT} = 2.5$ and $R_0^{NVT} = 2.0$. All other parameters as in the default set (see **Model parameterization in Supplementary Text**).

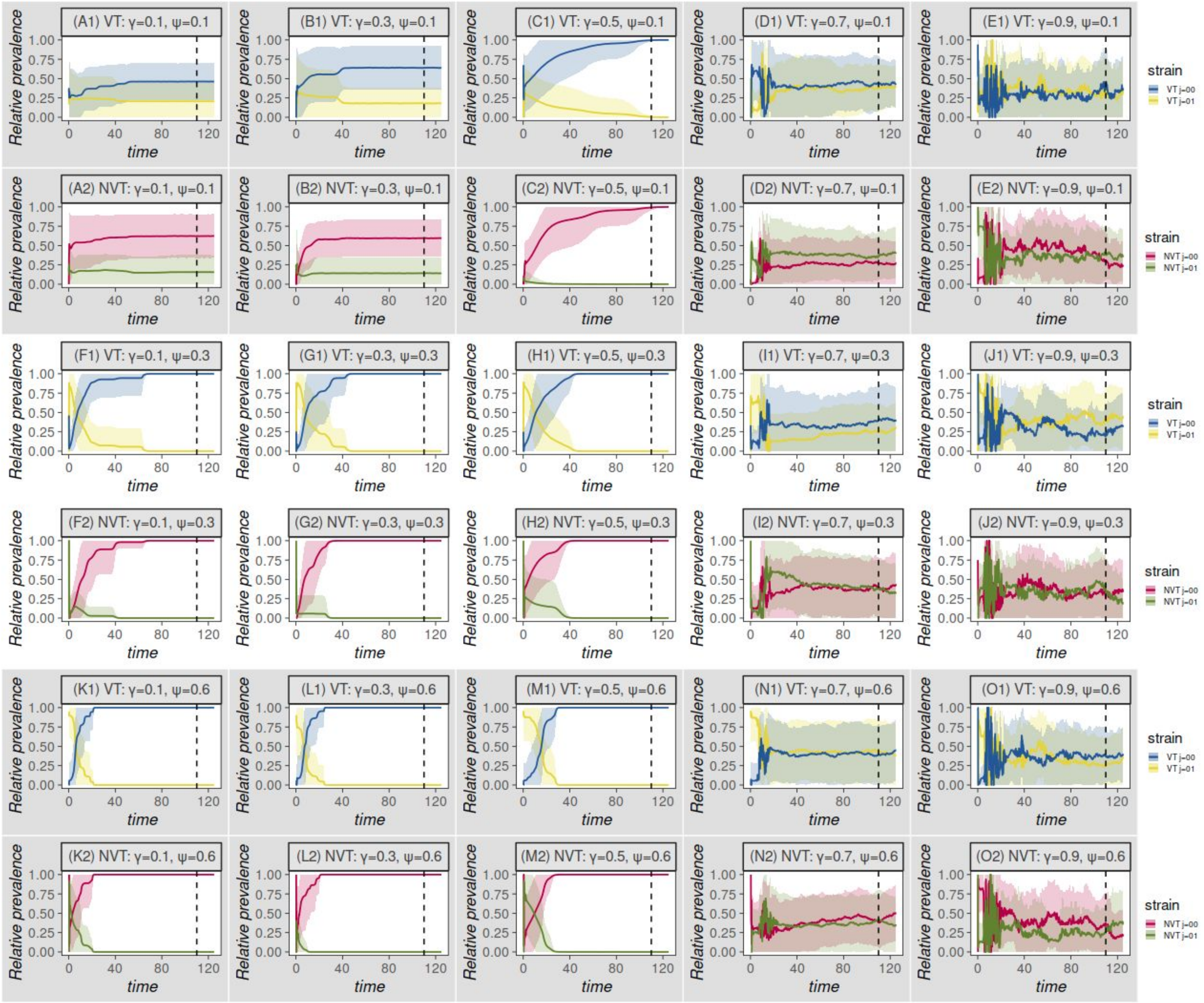


**Figure S6 - Examples of model output with no differences in R0 between VT and NVT.** Legend the same as in Supplementary Figure S5 but with no modelled differences in $R_0$, that is $R_0^{VT} = R_0^{NVT} = 2.5$.

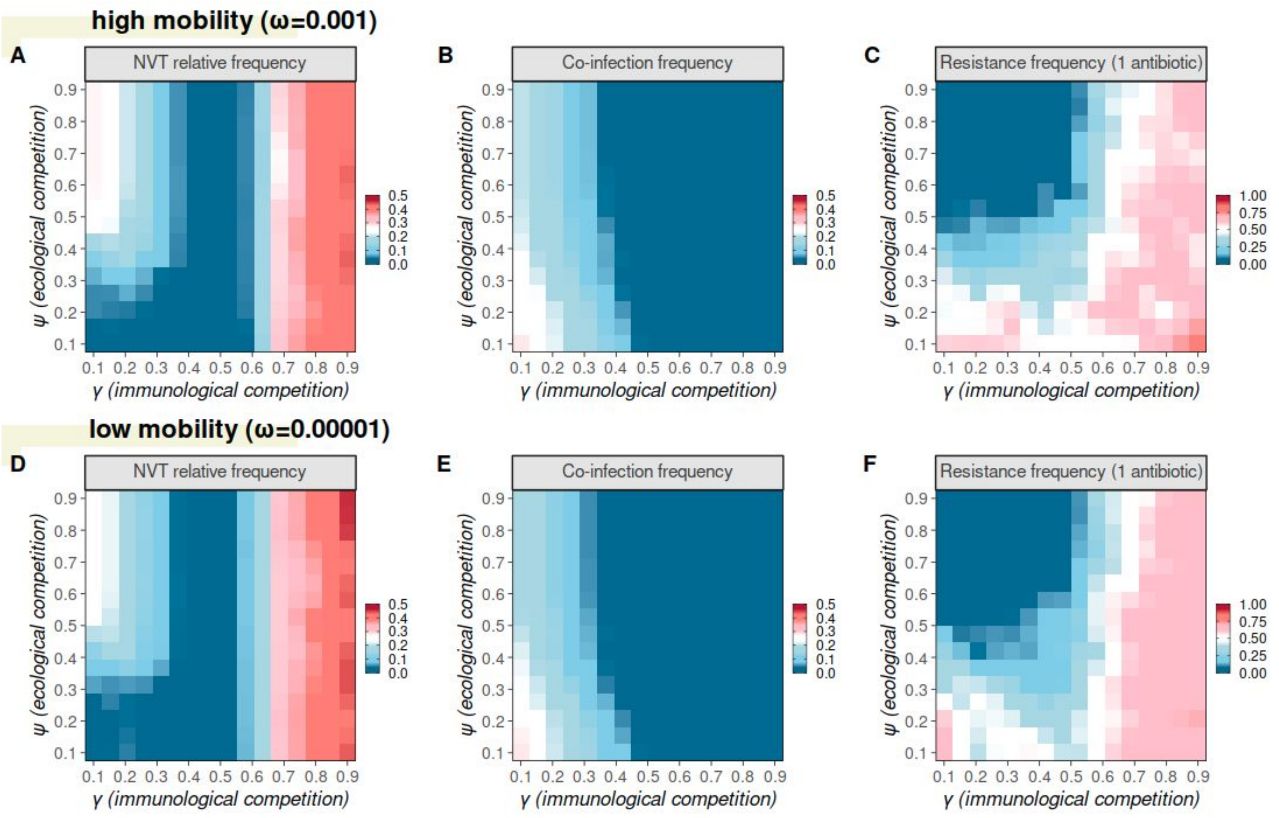


**Figure S7 - Model strain dynamics under variations to ecological and immunological competition under variations of mobility**. (A) Relative frequency (ratio) of total number of individuals carrying NVT versus carrying any type (NVT + VT). (B) Proportion of infected hosts carrying more than one strain (co-infection). (C) Relative frequency (ratio) of total number of individuals carrying resistant strains to one antibiotic (VT j=10, VT j=01, NVT j=10, NVT j=01) and carrying any strain. (A-C) All model parameters as in default parameter set except $\psi$, $\gamma$, varied in the y and x axis, respectively. Results presented are the mean over the last 5 years of a simulation, for particular combinations of $\psi$, $\gamma$. (A-F) Variation in mobility from $\omega = 0.001$ to $\omega = 0.00001$ as highlighted in the panel's titles.

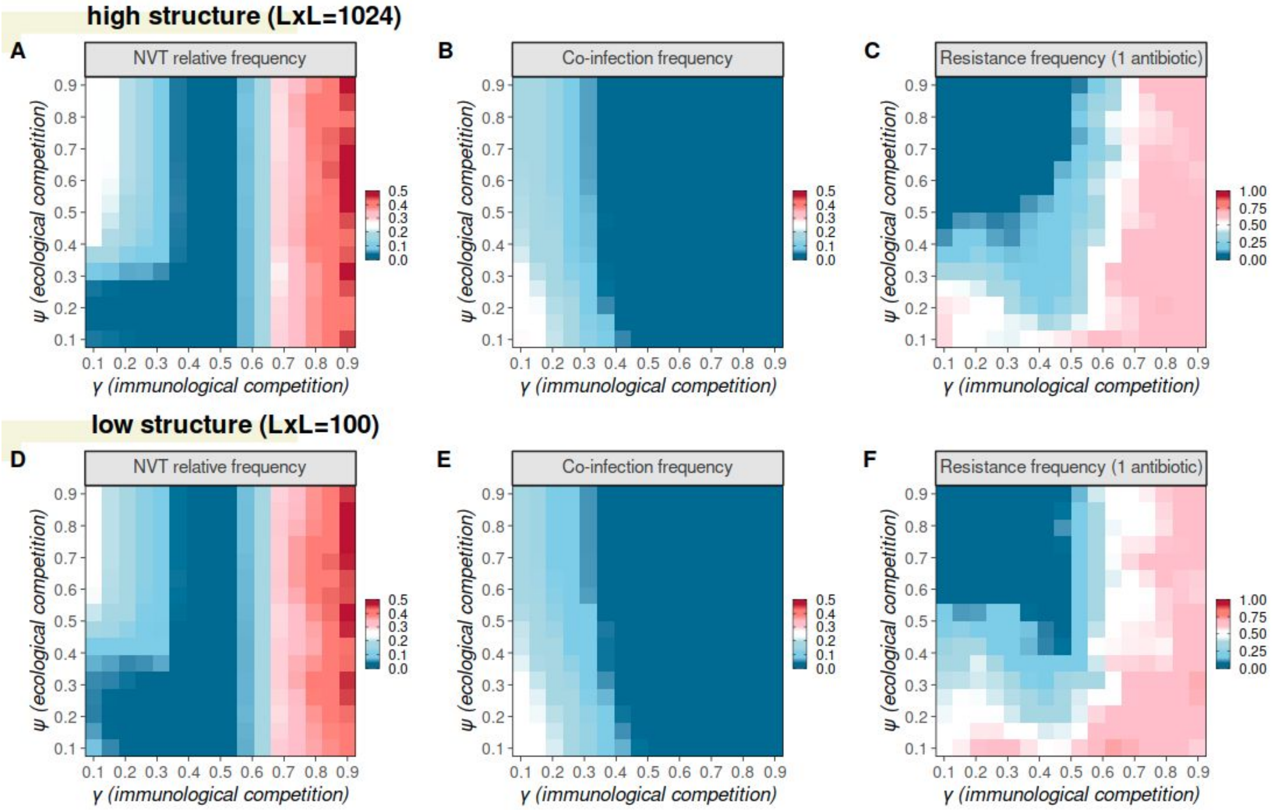


**Figure S8 - Model strain dynamics under variations to ecological and immunological competition under variations of population structure**. (A) Relative frequency (ratio) of total number of individuals carrying NVT versus carrying any type (NVT + VT). (B) Proportion of infected hosts carrying more than one strain (co-infection). (C) Relative frequency (ratio) of total number of individuals carrying resistant strains to one antibiotic (VT j=10, VT j=01, NVT j=10, NVT j=01) and carrying any strain. (A-C) All model parameters as in default parameter set except $\psi$, $\gamma$, varied in the y and x axis, respectively. Results presented are the mean over the last 5 years of a simulation, for particular combinations of $\psi$, $\gamma$. (A-F) Variation in mobility from LxL=100 to LxL=1024 as highlighted in the panel's titles.

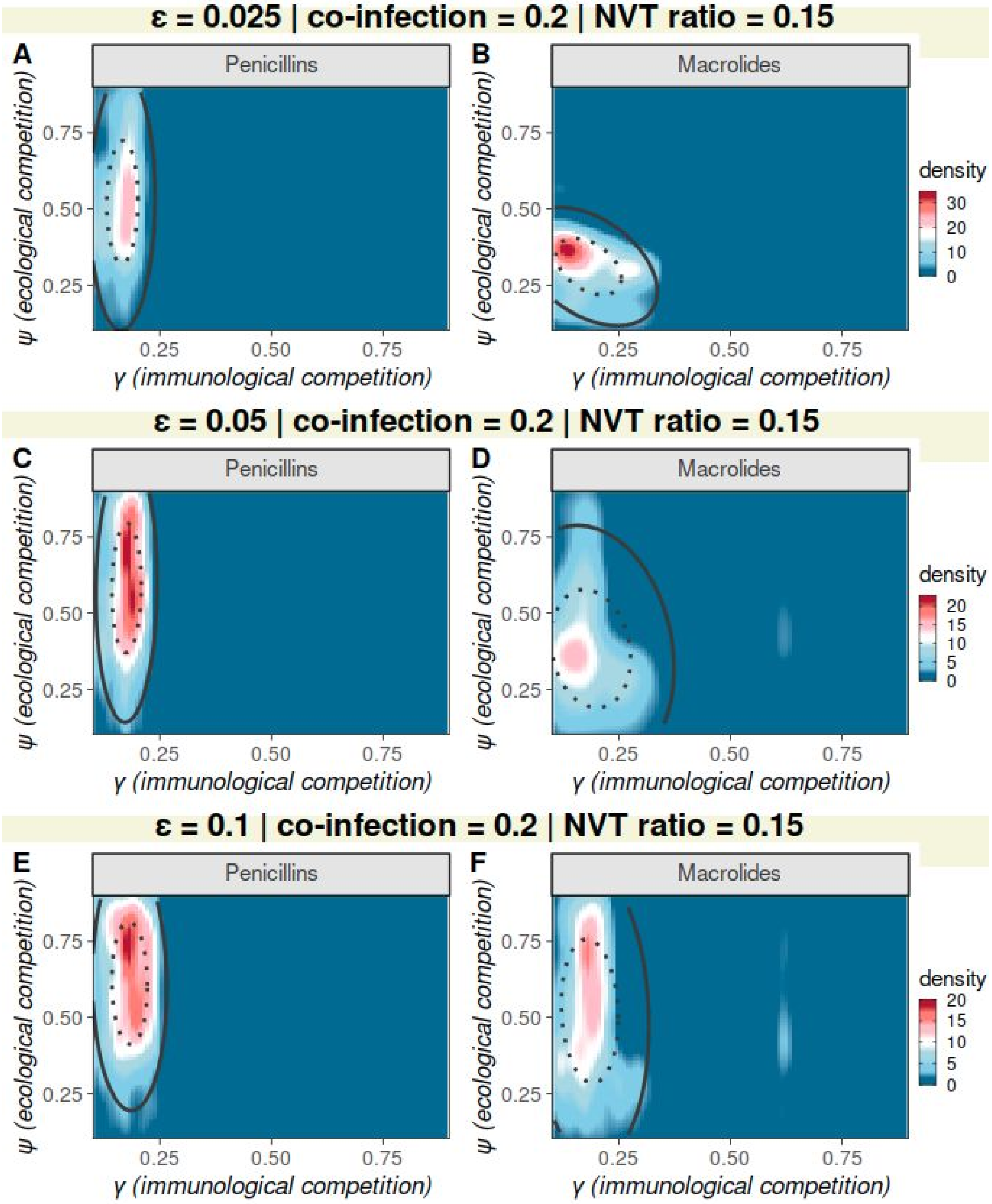


**Figure S9: Competition parameter space compatible with observed, pre-vaccination resistance levels when varying ABC sensitivity for co-infection target 0.2 and NVT ratio 0.15.** Approximate Bayesian Computation (ABC) output when varying immunological (γ) and ecological (ψ) competition, attempting to reproduce observed levels of resistance to penicillins (left column, A, C, E) and macrolides (right column, B, D, F) in the European region (see Data section in the main text for details). The colour scale is the density in ABC output of matches to observed resistance levels. Ellipses mark the 50 (dotted) and 95 (full) percentiles in ABC output. ABC priors and targets as detailed in the Methods section of the main text, the number of simulations was 6741, ε was varied at 0.025, 0.05, 0.1 as detailed in the panel's titles. The ABC targets of co-infection and NVT ratio were 0.2 and 0.15 respectively.

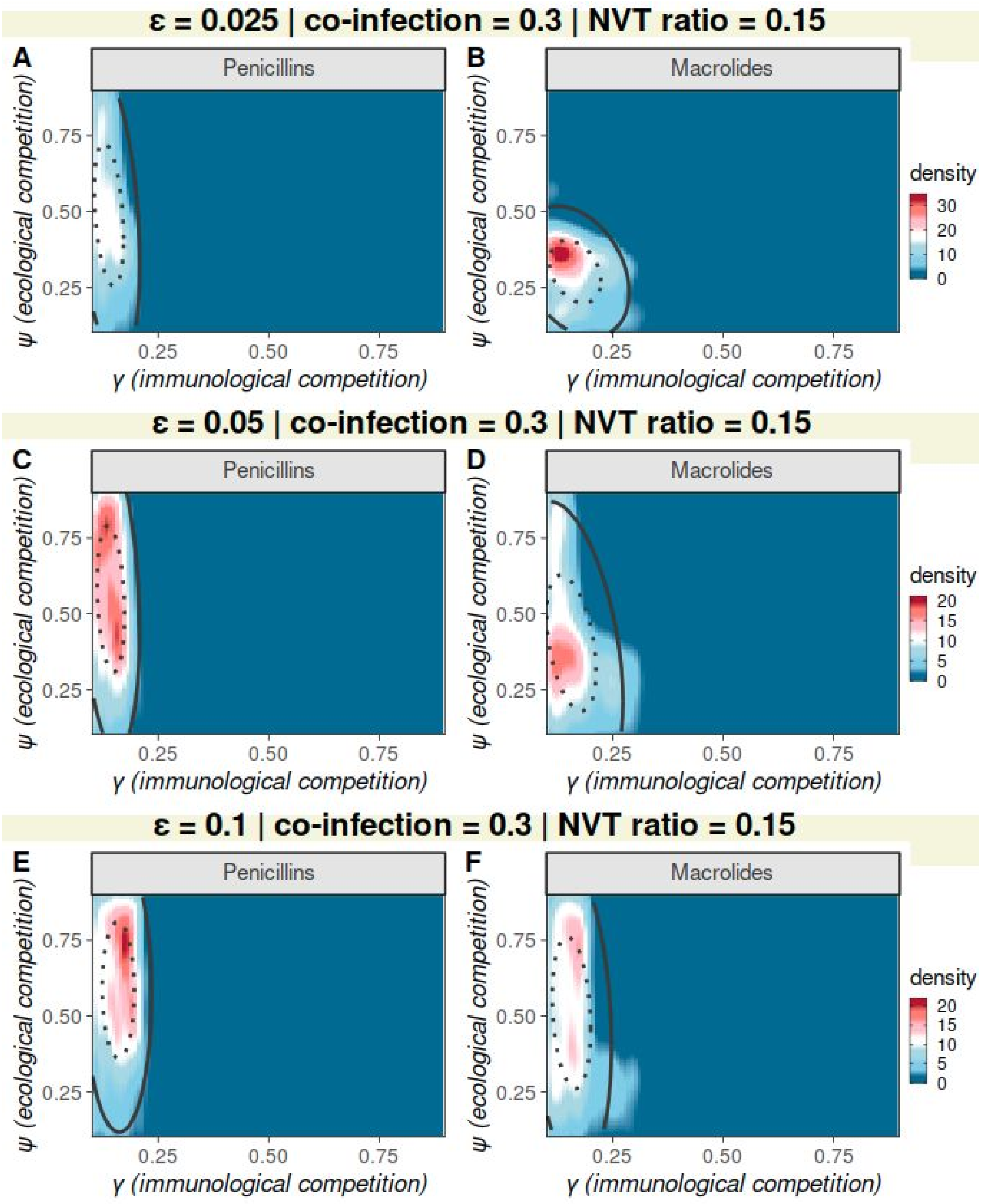


**Figure S10: Competition parameter space compatible with observed, pre-vaccination resistance levels when varying ABC sensitivity for co-infection target 0.3 and NVT ratio 0.15.** Approximate Bayesian Computation (ABC) output when varying immunological (γ) and ecological (ψ) competition, attempting to reproduce observed levels of resistance to penicillins (left column, A, C, E) and macrolides (right column, B, D, F) in the European region (see Data section in the main text for details). The colour scale is the density in ABC output of matches to observed resistance levels. Ellipses mark the 50 (dotted) and 95 (full) percentiles in ABC output. ABC priors and targets as detailed in the Methods section of the main text, the number of simulations was 6741, ε was varied at 0.025, 0.05, 0.1 as detailed in the panel's titles. The ABC targets of co-infection and NVT ratio were 0.3 and 0.15 respectively.
